## Supplementary Figures for "A high-resolution spatial map of cilia-associated proteins in the human fallopian tube"

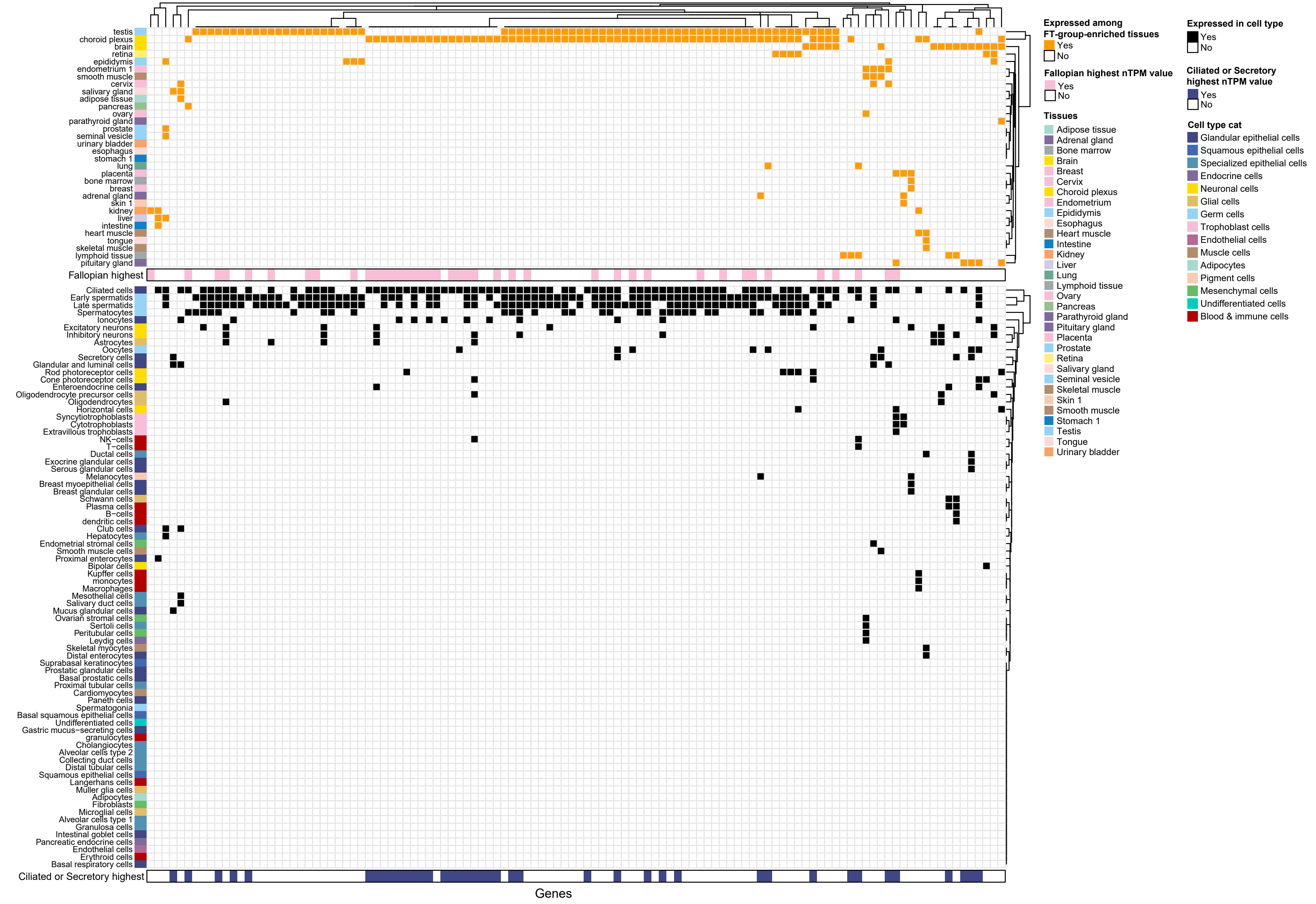
Expressed among FT-group-enriched tissues
Yes
No
Fallopian highest nTPM value
Yes
No
Tissues
Adipose tissue
Adrenal gland
Bone marrow
Brain
Breast
Cervix
Choroid plexus
Endometrium
Epididymis
Esophagus
Heart muscle
Intestine
Kidney
Liver
Lung
Lymphoid tissue
Ovary
Pancreas
Parathyroid gland
Pituitary gland
Placenta
Prostate
Retina
Salivary gland
Seminal vesicle
Skeletal muscle
Skin 1
Smooth muscle
Stomach 1
Testis
Tongue
Urinary bladder
Expressed in cell type
Yes
No
Ciliated or Secretory highest nTPM value
Yes
No
Cell type cat
Glandular epithelial cells
Squamous epithelial cells
Specialized epithelial cells
Endocrine cells
Neuronal cells
Glial cells
Germ cells
Trophoblast cells
Endothelial cells
Muscle cells
Adipocytes
Pigment cells
Mesenchymal cells
Undifferentiated cells
Blood & immune cells
testis
choroid plexus
brain
retina
epididymis
endometrium 1
smooth muscle
cervix
salivary gland
adipose tissue
pancreas
ovary
parathyroid gland
prostate
seminal vesicle
urinary bladder
esophagus
stomach 1
lung
placenta
bone marrow
breast
adrenal gland
skin 1
kidney
liver
intestine
heart muscle
tongue
skeletal muscle
lymphoid tissue
pituitary gland
Fallopian highest
Ciliated cells
Early spermatids
Late spermatids
Spermatocytes
Ionocytes
Excitatory neurons
Inhibitory neurons
Astrocytes
Oocytes
Secretory cells
Glandular and luminal cells
Rod photoreceptor cells
Cone photoreceptor cells
Enteroendocrine cells
Oligodendrocyte precursor cells
Oligodendrocytes
Horizontal cells
Syncytiotrophoblasts
Cytotrophoblasts
Extravillous trophoblasts
NK−cells
T−cells
Ductal cells
Exocrine glandular cells
Serous glandular cells
Melanocytes
Breast myoepithelial cells
Breast glandular cells
Schwann cells
Plasma cells
B−cells
dendritic cells
Club cells
Hepatocytes
Endometrial stromal cells
Smooth muscle cells
Proximal enterocytes
Bipolar cells
Kupffer cells
monocytes
Macrophages
Mesothelial cells
Salivary duct cells
Mucus glandular cells
Ovarian stromal cells
Sertoli cells
Peritubular cells
Leydig cells
Skeletal myocytes
Distal enterocytes
Suprabasal keratinocytes
Prostatic glandular cells
Basal prostatic cells
Proximal tubular cells
Cardiomyocytes
Paneth cells
Spermatogonia
Basal squamous epithelial cells
Undifferentiated cells
Gastric mucus−secreting cells
granulocytes
Cholangiocytes
Alveolar cells type 2
Collecting duct cells
Distal tubular cells
Squamous epithelial cells
Langerhans cells
Muller glia cells
Adipocytes
Fibroblasts
Microglial cells
Alveolar cells type 1
Granulosa cells
Intestinal goblet cells
Pancreatic endocrine cells
Endothelial cells
Erythroid cells
Basal respiratory cells
Ciliated or Secretory highest
Genes

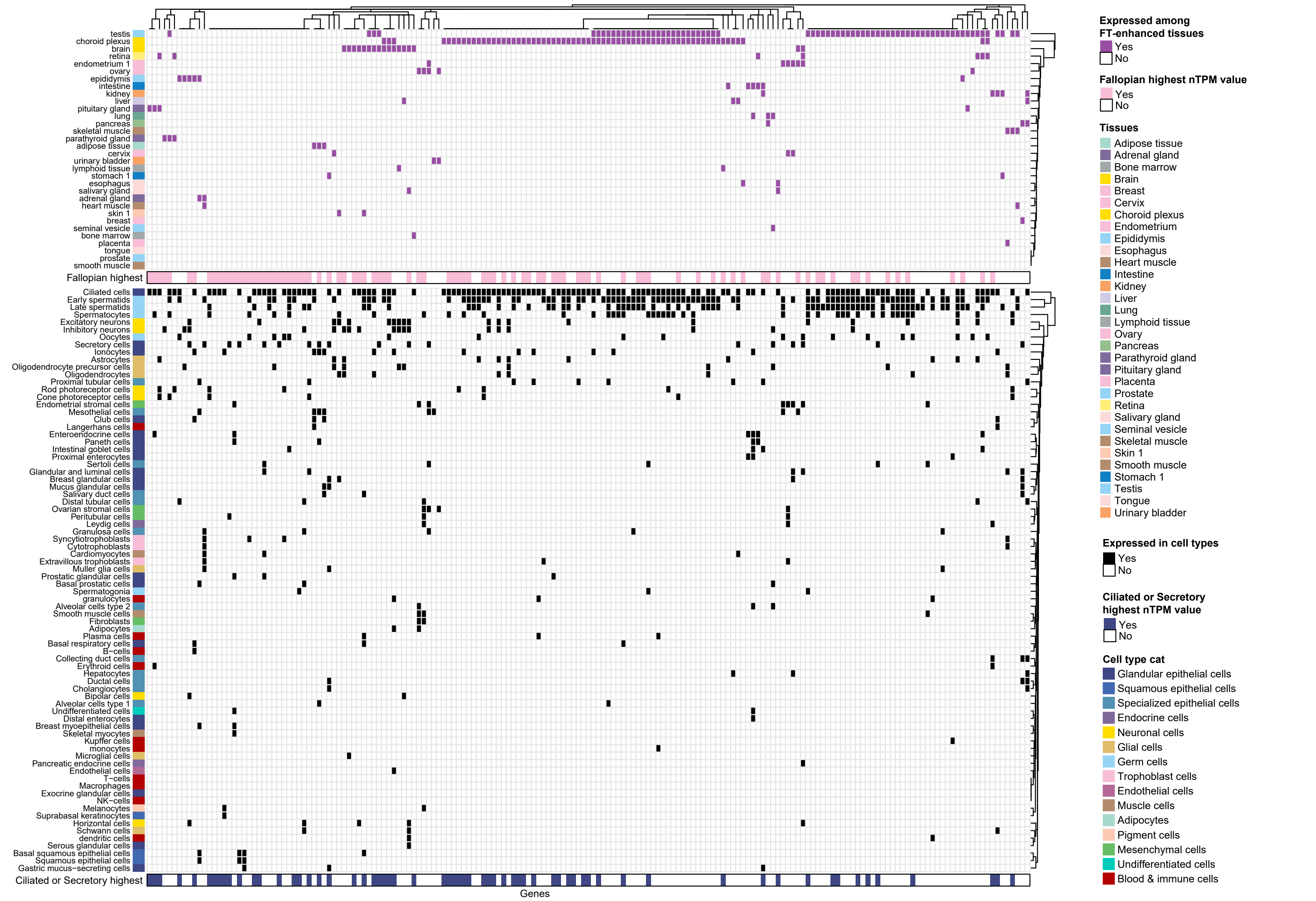
testis
choroid plexus
brain
retina
endometrium 1
ovary
epididymis
intestine
kidney
liver
pituitary gland
lung
pancreas
skeletal muscle
parathyroid gland
adipose tissue
cervix
urinary bladder
lymphoid tissue
stomach 1
esophagus
salivary gland
adrenal gland
heart muscle
skin 1
breast
seminal vesicle
bone marrow
placenta
tongue
prostate
smooth muscle
Fallopian highest
Ciliated cells
Early spermatids
Late spermatids
Spermatocytes
Excitatory neurons
Inhibitory neurons
Oocytes
Secretory cells
Ionocytes
Astrocytes
Oligodendrocyte precursor cells
Oligodendrocytes
Proximal tubular cells
Rod photoreceptor cells
Cone photoreceptor cells
Endometrial stromal cells
Mesothelial cells
Club cells
Langerhans cells
Enteroendocrine cells
Paneth cells
Intestinal goblet cells
Proximal enterocytes
Sertoli cells
Glandular and luminal cells
Breast glandular cells
Mucus glandular cells
Salivary duct cells
Distal tubular cells
Ovarian stromal cells
Peritubular cells
Leydig cells
Granulosa cells
Syncytiotrophoblasts
Cytotrophoblasts
Cardiomyocytes
Extravillous trophoblasts
Muller glia cells
Prostatic glandular cells
Basal prostatic cells
Spermatogonia
granulocytes
Alveolar cells type 2
Smooth muscle cells
Fibroblasts
Adipocytes
Plasma cells
Basal respiratory cells
B−cells
Collecting duct cells
Erythroid cells
Hepatocytes
Ductal cells
Cholangiocytes
Bipolar cells
Alveolar cells type 1
Undifferentiated cells
Distal enterocytes
Breast myoepithelial cells
Skeletal myocytes
Kupffer cells
monocytes
Microglial cells
Pancreatic endocrine cells
Endothelial cells
T−cells
Macrophages
Exocrine glandular cells
NK−cells
Melanocytes
Suprabasal keratinocytes
Horizontal cells
Schwann cells
dendritic cells
Serous glandular cells
Basal squamous epithelial cells
Squamous epithelial cells
Gastric mucus−secreting cells
Ciliated or Secretory highest
Genes
Expressed among FT-enhanced tissues
Yes
No
Fallopian highest nTPM value
Yes
No
Tissues
Adipose tissue
Adrenal gland
Bone marrow
Brain
Breast
Cervix
Choroid plexus
Endometrium
Epididymis
Esophagus
Heart muscle
Intestine
Kidney
Liver
Lung
Lymphoid tissue
Ovary
Pancreas
Parathyroid gland
Pituitary gland
Placenta
Prostate
Retina
Salivary gland
Seminal vesicle
Skeletal muscle
Skin 1
Smooth muscle
Stomach 1
Testis
Tongue
Urinary bladder
Expressed in cell types
Yes
No
Ciliated or Secretory highest nTPM value
Yes
No
Cell type cat
Glandular epithelial cells
Squamous epithelial cells
Specialized epithelial cells
Endocrine cells
Neuronal cells
Glial cells
Germ cells
Trophoblast cells
Endothelial cells
Muscle cells
Adipocytes
Pigment cells
Mesenchymal cells
Undifferentiated cells
Blood & immune cells

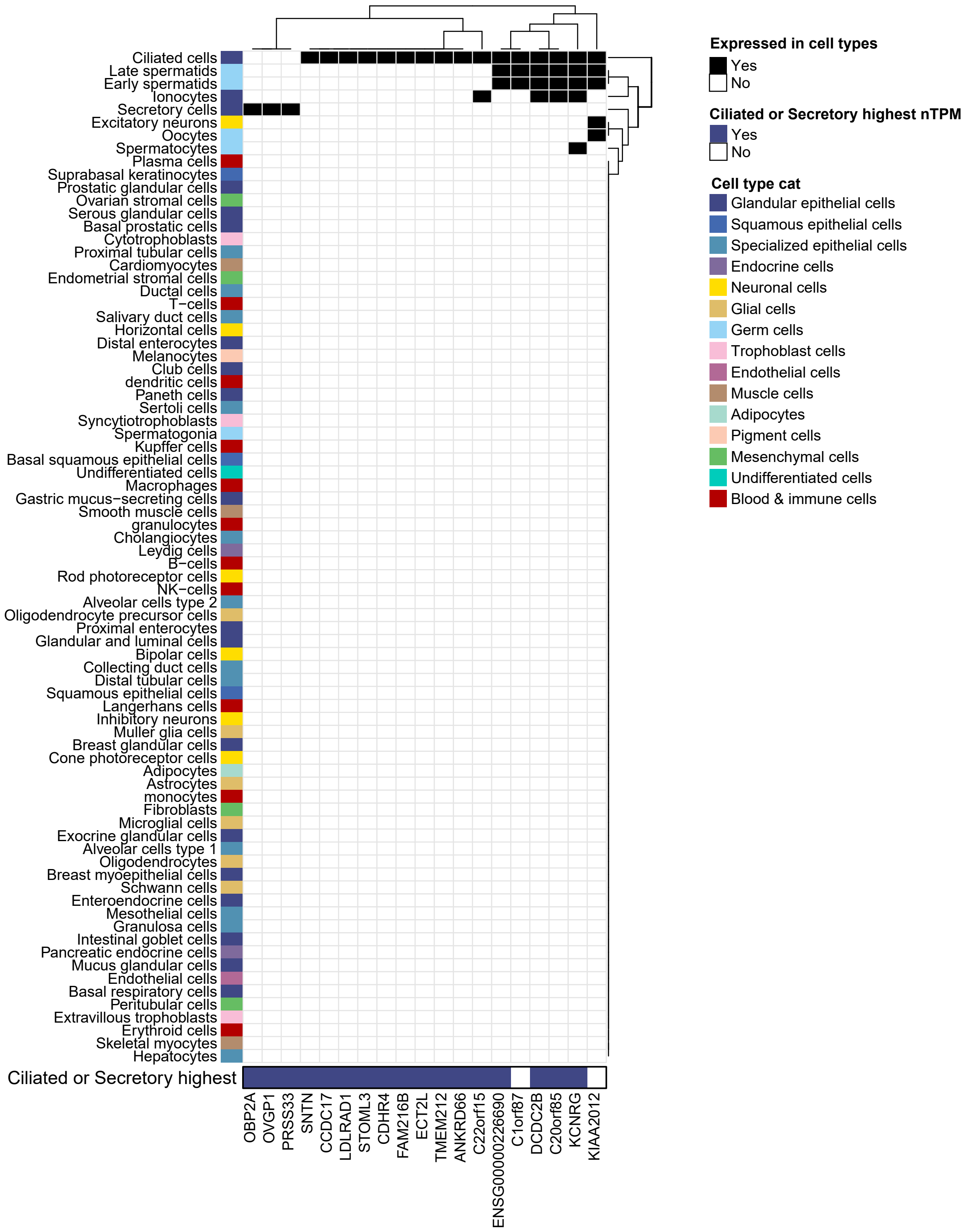

Expressed in cell types
Yes
No
Ciliated or Secretory highest nTPM
Yes
No
Cell type cat
Glandular epithelial cells
Squamous epithelial cells
Specialized epithelial cells
Endocrine cells
Neuronal cells
Glial cells
Germ cells
Trophoblast cells
Endothelial cells
Muscle cells
Adipocytes
Pigment cells
Mesenchymal cells
Undifferentiated cells
Blood & immune cells
Ciliated cells
Late spermatids
Early spermatids
Ionocytes
Secretory cells
Excitatory neurons
Oocytes
Spermatocytes
Plasma cells
Suprabasal keratinocytes
Prostatic glandular cells
Ovarian stromal cells
Serous glandular cells
Basal prostatic cells
Cytotrophoblasts
Proximal tubular cells
Cardiomyocytes
Endometrial stromal cells
Ductal cells
T−cells
Salivary duct cells
Horizontal cells
Distal enterocytes
Melanocytes
Club cells
dendritic cells
Paneth cells
Sertoli cells
Syncytiotrophoblasts
Spermatogonia
Kupffer cells
Basal squamous epithelial cells
Undifferentiated cells
Macrophages
Gastric mucus−secreting cells
Smooth muscle cells
granulocytes
Cholangiocytes
Leydig cells
B−cells
Rod photoreceptor cells
NK−cells
Alveolar cells type 2
Oligodendrocyte precursor cells
Proximal enterocytes
Glandular and luminal cells
Bipolar cells
Collecting duct cells
Distal tubular cells
Squamous epithelial cells
Langerhans cells
Inhibitory neurons
Muller glia cells
Breast glandular cells
Cone photoreceptor cells
Adipocytes
Astrocytes
monocytes
Fibroblasts
Microglial cells
Exocrine glandular cells
Alveolar cells type 1
Oligodendrocytes
Breast myoepithelial cells
Schwann cells
Enteroendocrine cells
Mesothelial cells
Granulosa cells
Intestinal goblet cells
Pancreatic endocrine cells
Mucus glandular cells
Endothelial cells
Basal respiratory cells
Peritubular cells
Extravillous trophoblasts
Erythroid cells
Skeletal myocytes
Hepatocytes
Ciliated or Secretory highest
OBP2A
OVGP1
PRSS33
SNTN
CCDC17
LDLRAD1
STOML3
CDHR4
FAM216B
ECT2L
TMEM212
ANKRD66
C22orf15
ENSG00000226690
C1orf87
DCDC2B
C20orf85
KCNRG
KIAA2012

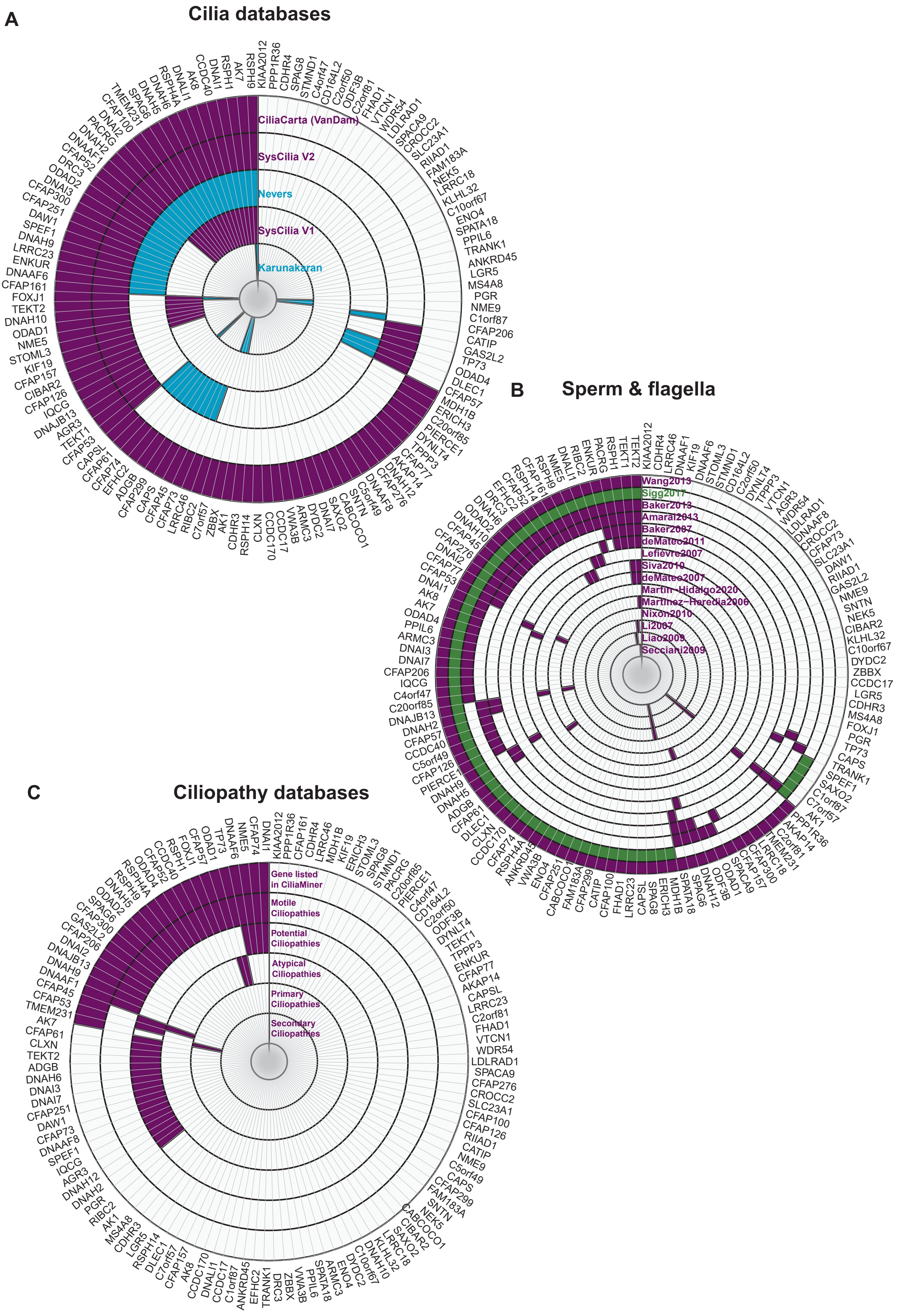
A
Cilia databases
CiliaCarta (VanDam)
SysCilia V2
Nevers
SysCilia V1
Karunakaran
B
Sperm & flagella
Wang2013
Sigg2017
Baker2013
Amaral2013
Baker2007
deMateo2011
Lefièvre2007
Siva2010
deMateo2007
Martin-Hidalgo2020
Martinez-Heredia2006
Nixon2010
Li2007
Liao2009
Secciani2009
C
Ciliopathy databases
Gene listed in CiliaMiner
Motile Ciliopathies
Potential Ciliopathies
Atypical Ciliopathies
Primary Ciliopathies
Secondary Ciliopathies

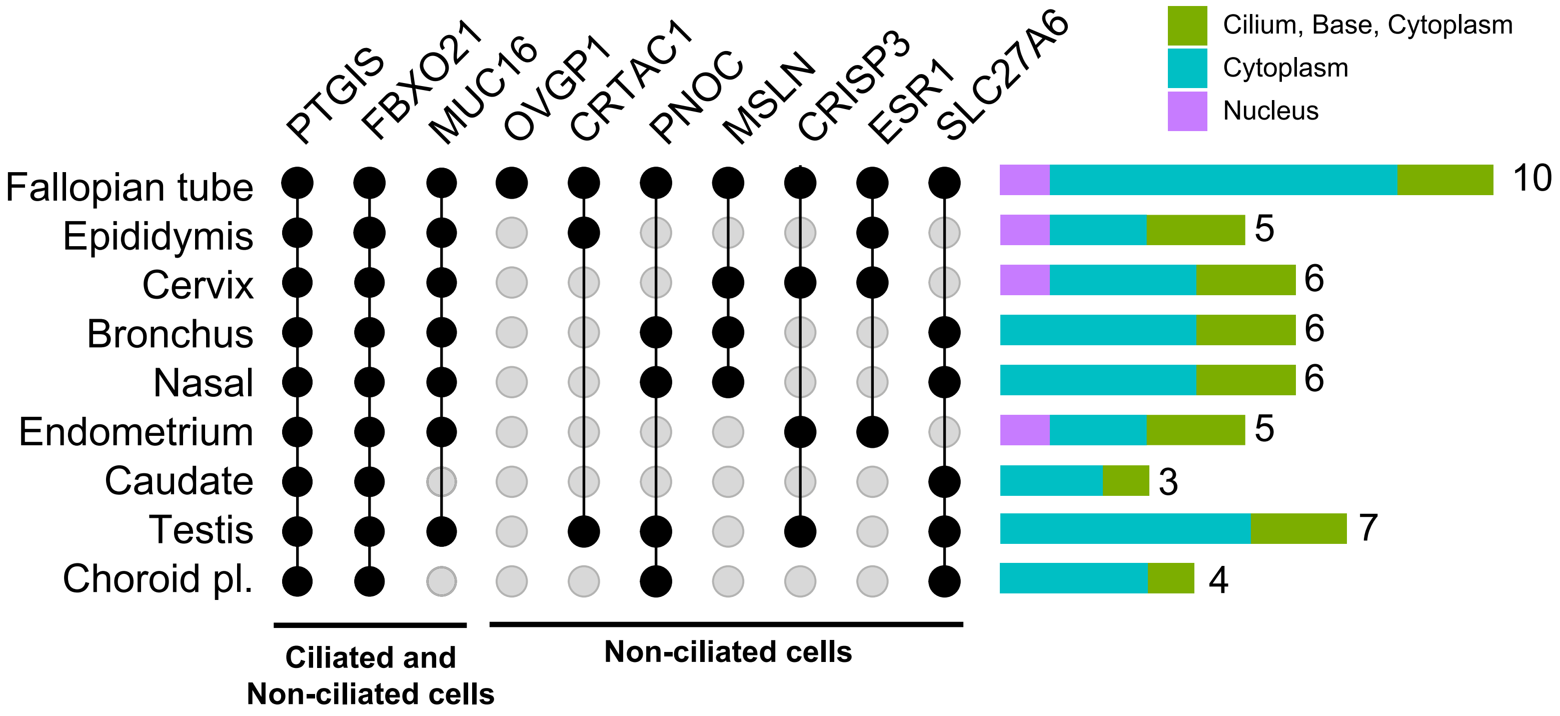

PTGIS
FBXO21
MUC16
OVGP1
CRTAC1
PNOC
MSLN
CRISP3
ESR1
SLC27A6
Cilium, Base, Cytoplasm
Cytoplasm
Nucleus
Fallopian tube
Epididymis
Cervix
Bronchus
Nasal
Endometrium
Caudate
Testis
Choroid pl.
10
5
6
6
6
5
3
7
4
Ciliated and Non-ciliated cells
Non-ciliated cells

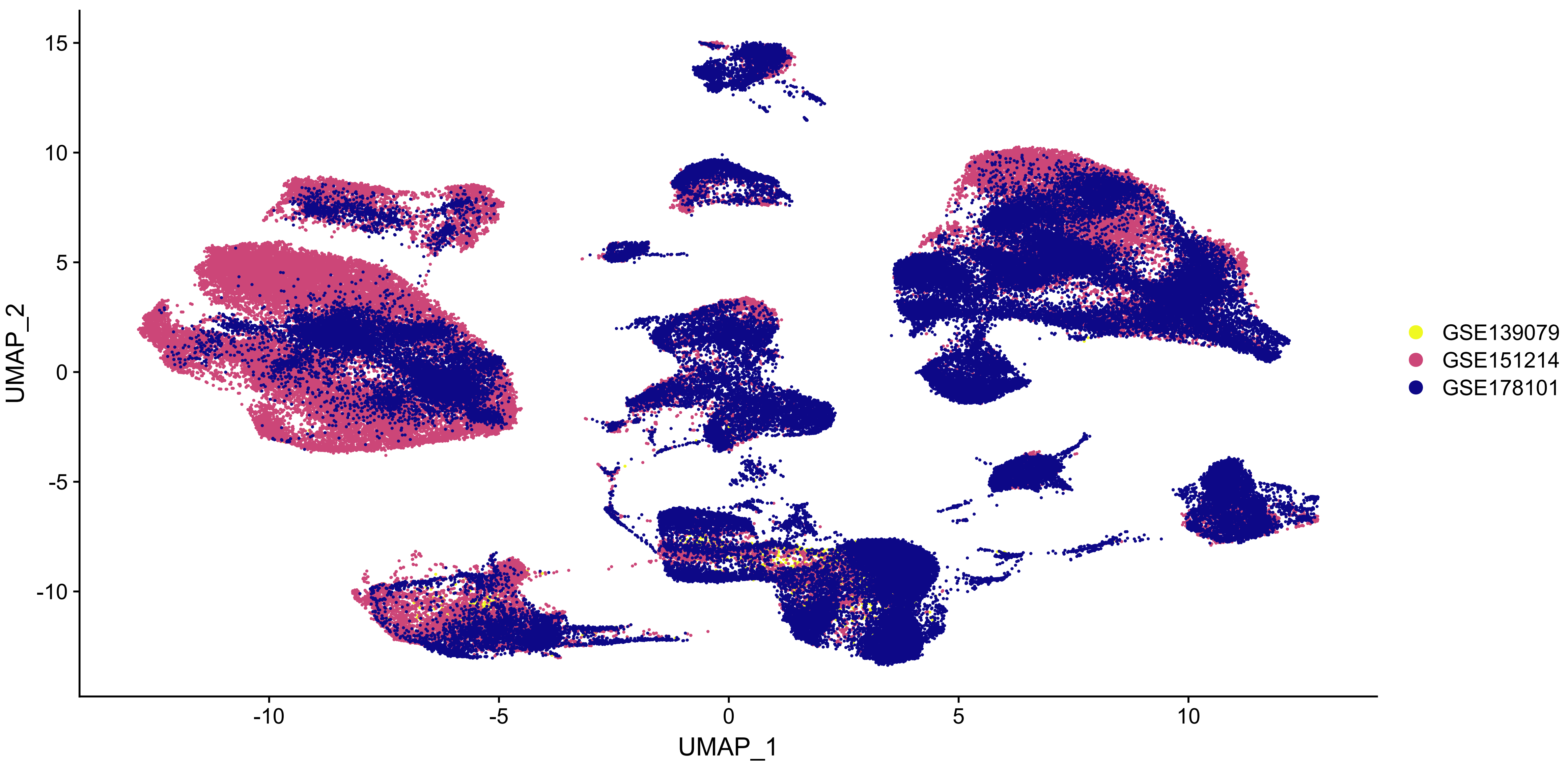

UMAP_2
UMAP_1
GSE139079
GSE151214
GSE178101


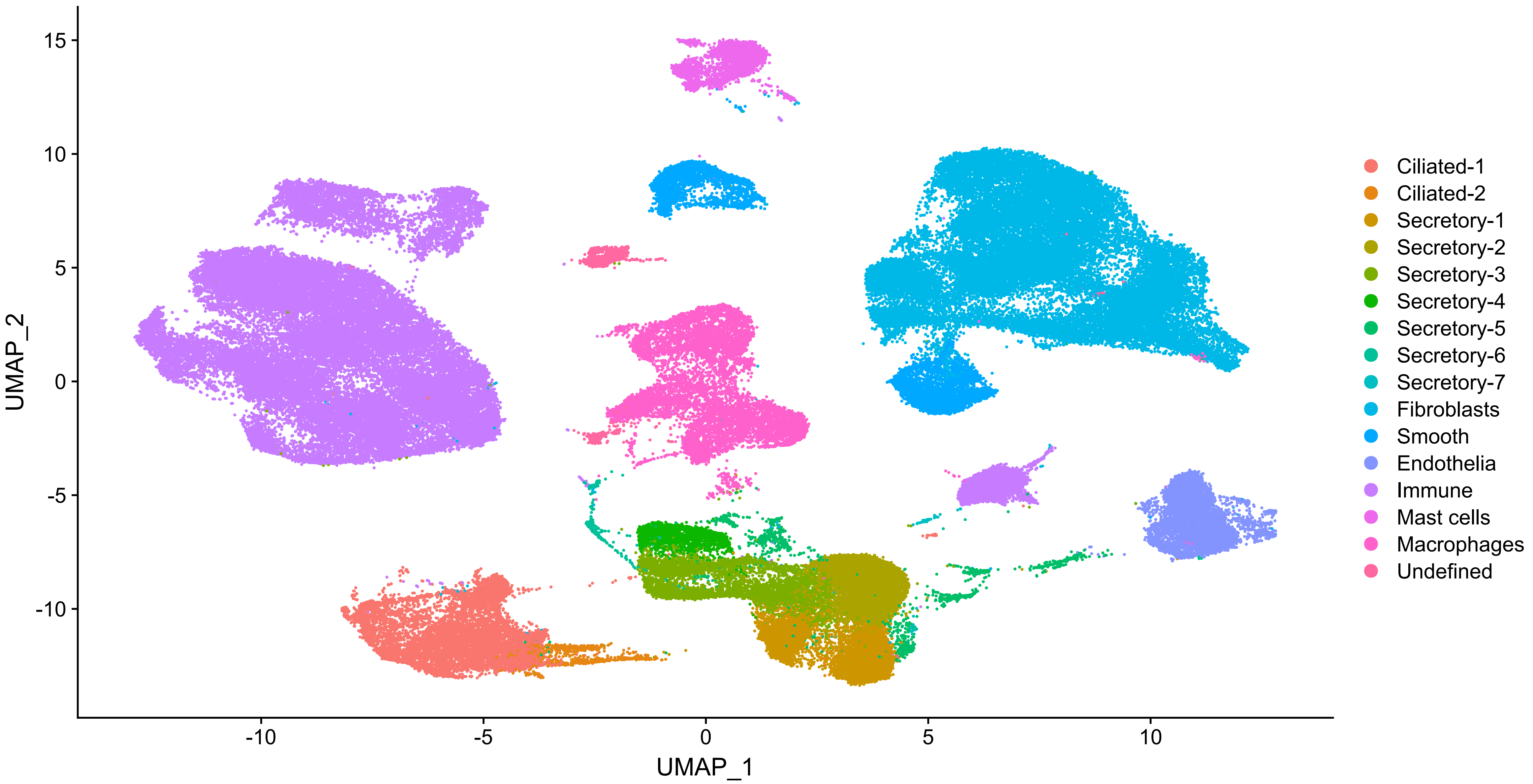

UMAP_2
UMAP_1
Ciliated-1
Ciliated-2
Secretory-1
Secretory-2
Secretory-3
Secretory-4
Secretory-5
Secretory-6
Secretory-7
Fibroblasts
Smooth
Endothelia
Immune
Mast cells
Macrophages
Undefined

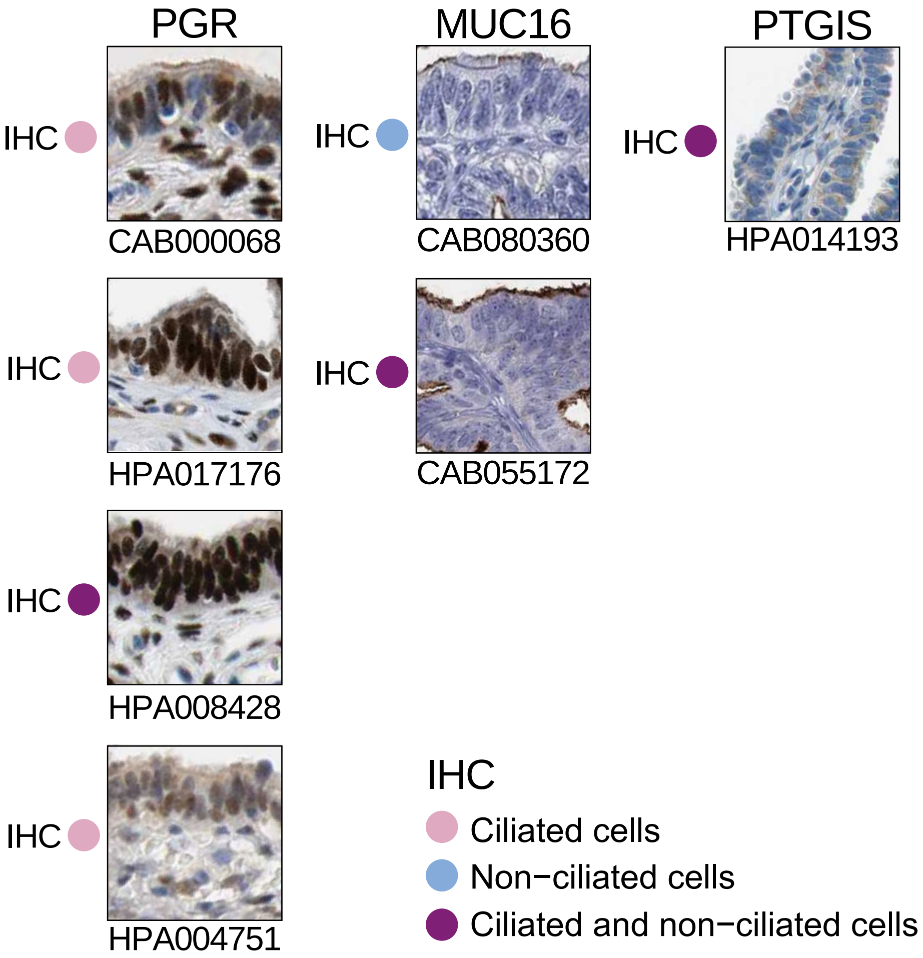
PGR
MUC16
PTGIS
IHC
CAB000068
CAB080360
HPA014193
IHC
HPA017176
CAB055172
IHC
HPA008428
IHC
HPA004751
IHC
Ciliated cells
Non−ciliated cells
Ciliated and non−ciliated cells

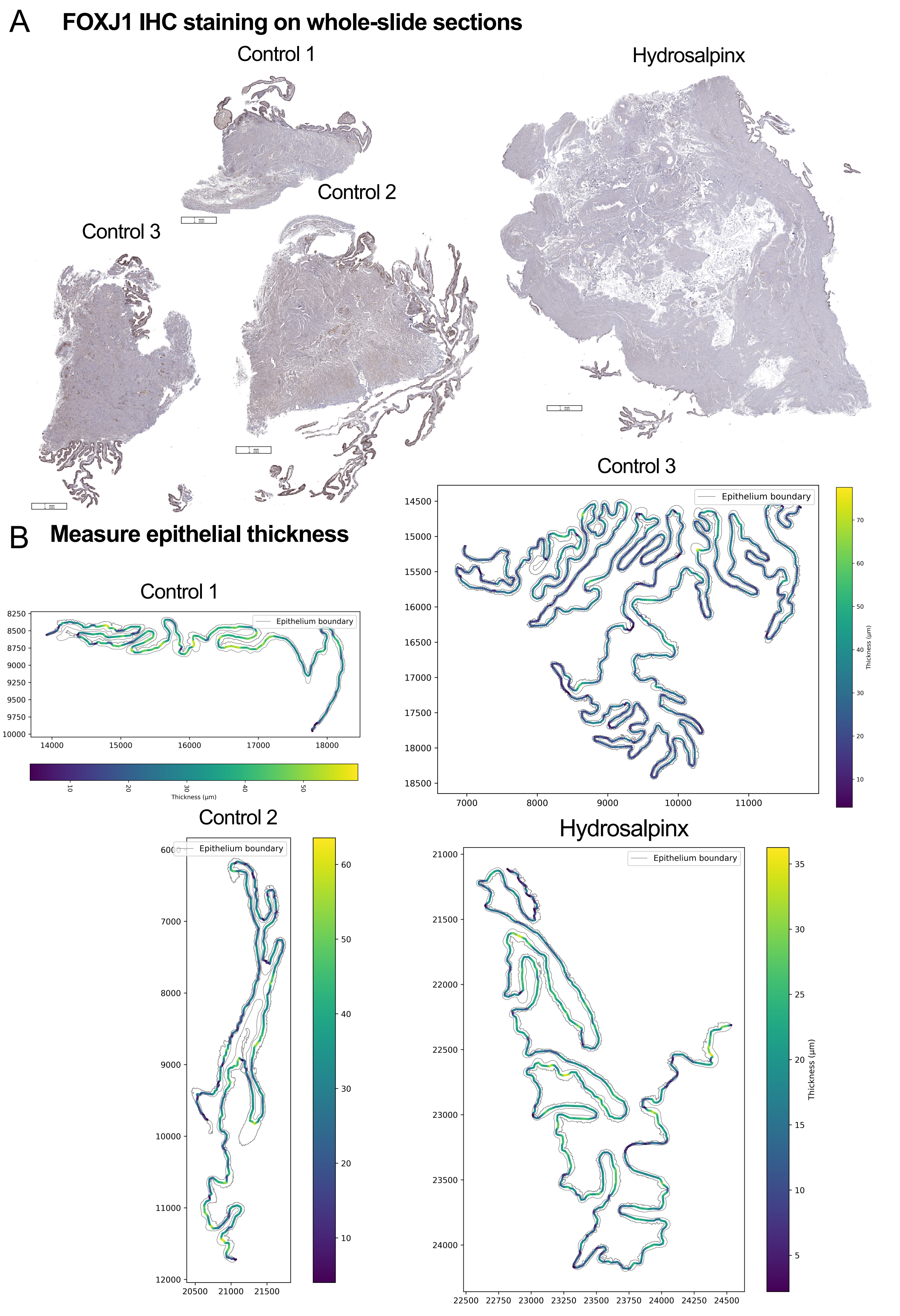
A
FOXJ1 IHC staining on whole-slide sections
Control 1
Hydrosalpinx
Control 2
Control 3
1 mm
B
Measure epithelial thickness
Control 1
Epithelium boundary
Thickness (μm)
Control 3
Epithelium boundary
Thickness (μm)
Control 2
Epithelium boundary
Hydrosalpinx
Epithelium boundary
Thickness (μm)

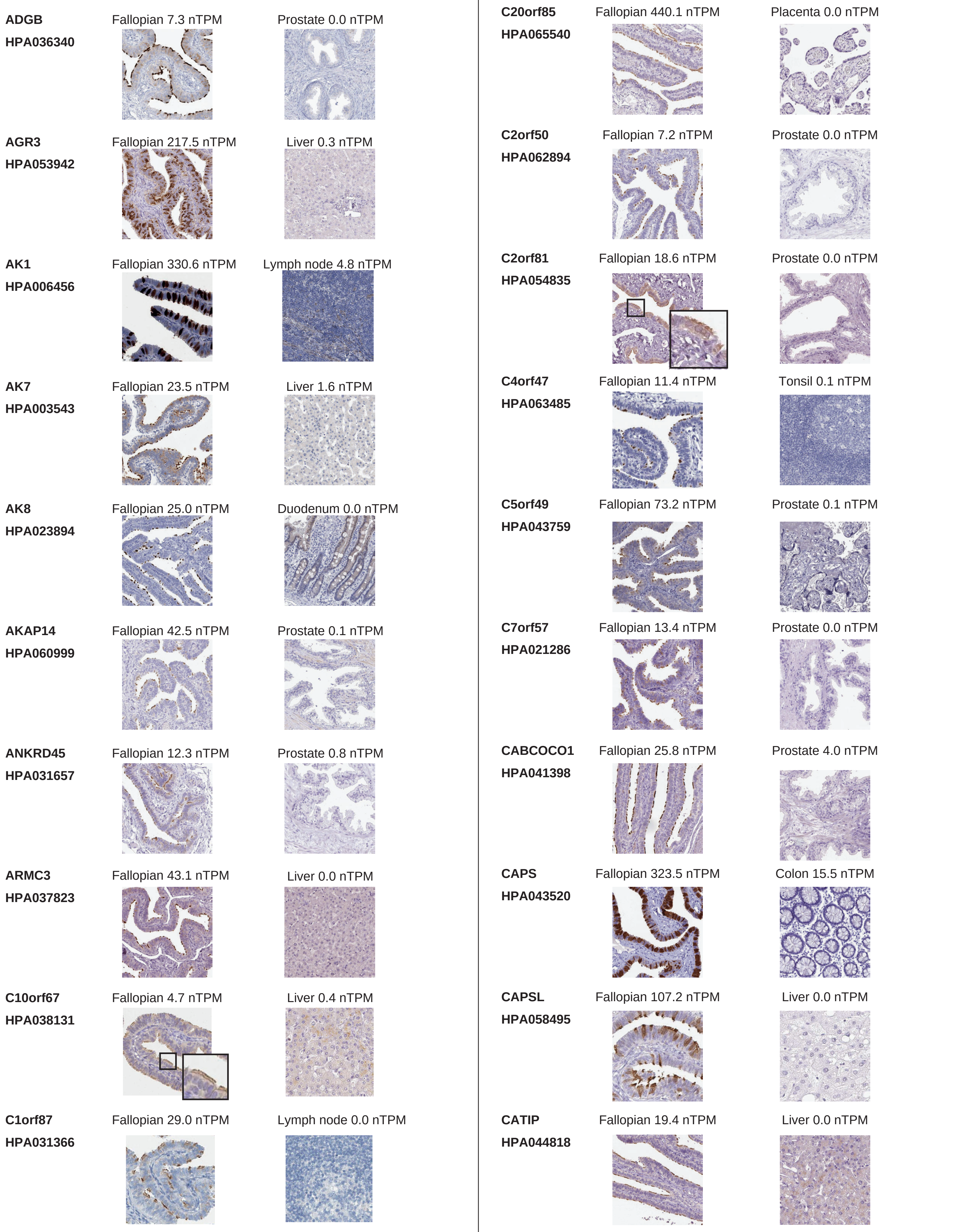
ADGB
HPA036340
Fallopian 7.3 nTPM
Prostate 0.0 nTPM
AGR3
HPA053942
Fallopian 217.5 nTPM
Liver 0.3 nTPM
AK1
HPA006456
Fallopian 330.6 nTPM
Lymph node 4.8 nTPM
AK7
HPA003543
Fallopian 23.5 nTPM
Liver 1.6 nTPM
AK8
HPA023894
Fallopian 25.0 nTPM
Duodenum 0.0 nTPM
AKAP14
HPA060999
Fallopian 42.5 nTPM
Prostate 0.1 nTPM
ANKRD45
HPA031657
Fallopian 12.3 nTPM
Prostate 0.8 nTPM
ARMC3
HPA037823
Fallopian 43.1 nTPM
Liver 0.0 nTPM
C10orf67
HPA038131
Fallopian 4.7 nTPM
Liver 0.4 nTPM
C1orf87
HPA031366
Fallopian 29.0 nTPM
Lymph node 0.0 nTPM
C20orf85
HPA065540
Fallopian 440.1 nTPM
Placenta 0.0 nTPM
C2orf50
HPA062894
Fallopian 7.2 nTPM
Prostate 0.0 nTPM
C2orf81
HPA054835
Fallopian 18.6 nTPM
Prostate 0.0 nTPM
C4orf47
HPA063485
Fallopian 11.4 nTPM
Tonsil 0.1 nTPM
C5orf49
HPA043759
Fallopian 73.2 nTPM
Prostate 0.1 nTPM
C7orf57
HPA021286
Fallopian 13.4 nTPM
Prostate 0.0 nTPM
CABCOCO1
HPA041398
Fallopian 25.8 nTPM
Prostate 4.0 nTPM
CAPS
HPA043520
Fallopian 323.5 nTPM
Colon 15.5 nTPM
CAPSL
HPA058495
Fallopian 107.2 nTPM
Liver 0.0 nTPM
CATIP
HPA044818
Fallopian 19.4 nTPM
Liver 0.0 nTPM

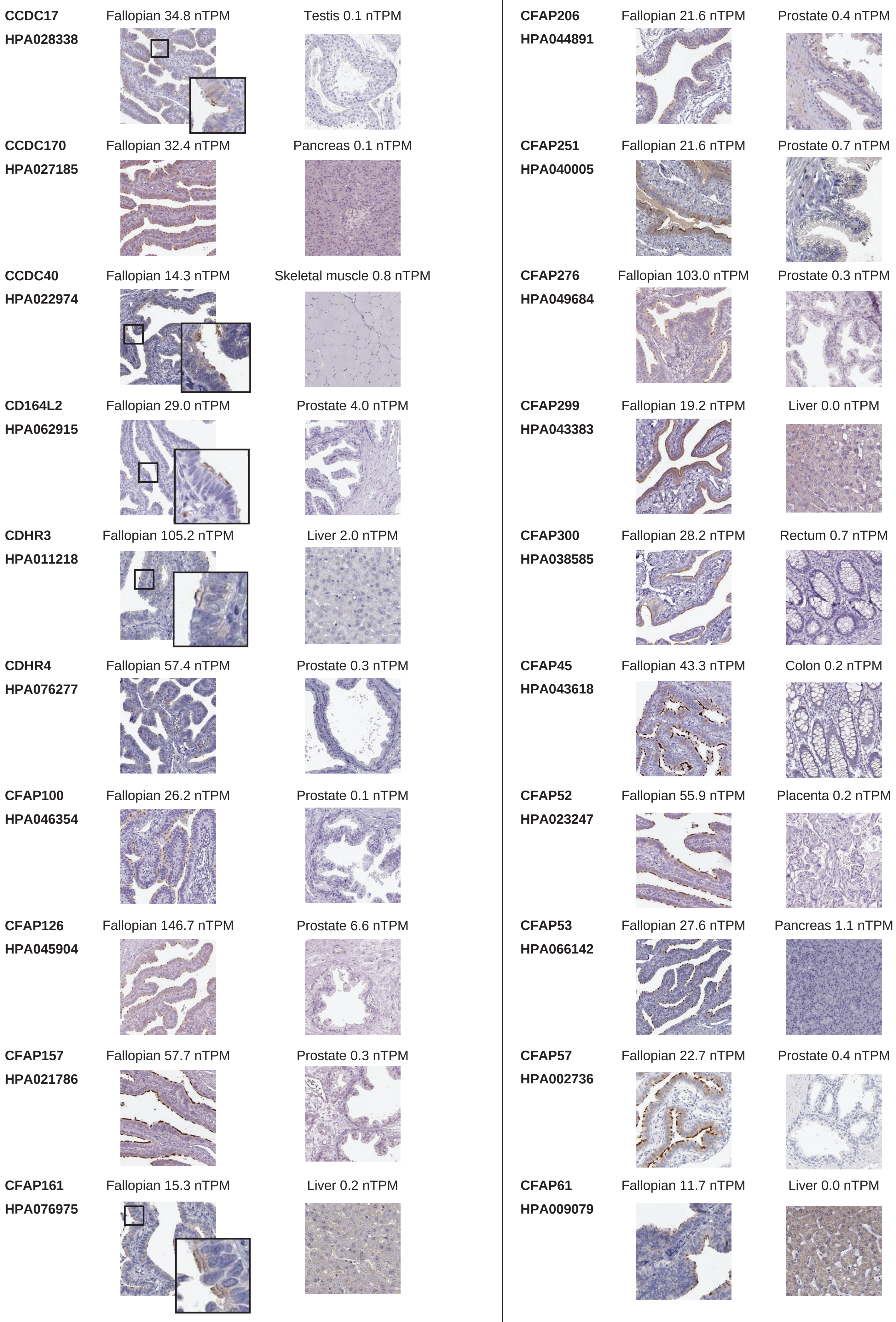
CCDC17
HPA028338
Fallopian 34.8 nTPM
Testis 0.1 nTPM
CCDC170
HPA027185
Fallopian 32.4 nTPM
Pancreas 0.1 nTPM
CCDC40
HPA022974
Fallopian 14.3 nTPM
Skeletal muscle 0.8 nTPM
CD164L2
HPA062915
Fallopian 29.0 nTPM
Prostate 4.0 nTPM
CDHR3
HPA011218
Fallopian 105.2 nTPM
Liver 2.0 nTPM
CDHR4
HPA076277
Fallopian 57.4 nTPM
Prostate 0.3 nTPM
CFAP100
HPA046354
Fallopian 26.2 nTPM
Prostate 0.1 nTPM
CFAP126
HPA045904
Fallopian 146.7 nTPM
Prostate 6.6 nTPM
CFAP157
HPA021786
Fallopian 57.7 nTPM
Prostate 0.3 nTPM
CFAP161
HPA076975
Fallopian 15.3 nTPM
Liver 0.2 nTPM
CFAP206
HPA044891
Fallopian 21.6 nTPM
Prostate 0.4 nTPM
CFAP251
HPA040005
Fallopian 21.6 nTPM
Prostate 0.7 nTPM
CFAP276
HPA049684
Fallopian 103.0 nTPM
Prostate 0.3 nTPM
CFAP299
HPA043383
Fallopian 19.2 nTPM
Liver 0.0 nTPM
CFAP300
HPA038585
Fallopian 28.2 nTPM
Rectum 0.7 nTPM
CFAP45
HPA043618
Fallopian 43.3 nTPM
Colon 0.2 nTPM
CFAP52
HPA023247
Fallopian 55.9 nTPM
Placenta 0.2 nTPM
CFAP53
HPA066142
Fallopian 27.6 nTPM
Pancreas 1.1 nTPM
CFAP57
HPA002736
Fallopian 22.7 nTPM
Prostate 0.4 nTPM
CFAP61
HPA009079
Fallopian 11.7 nTPM
Liver 0.0 nTPM

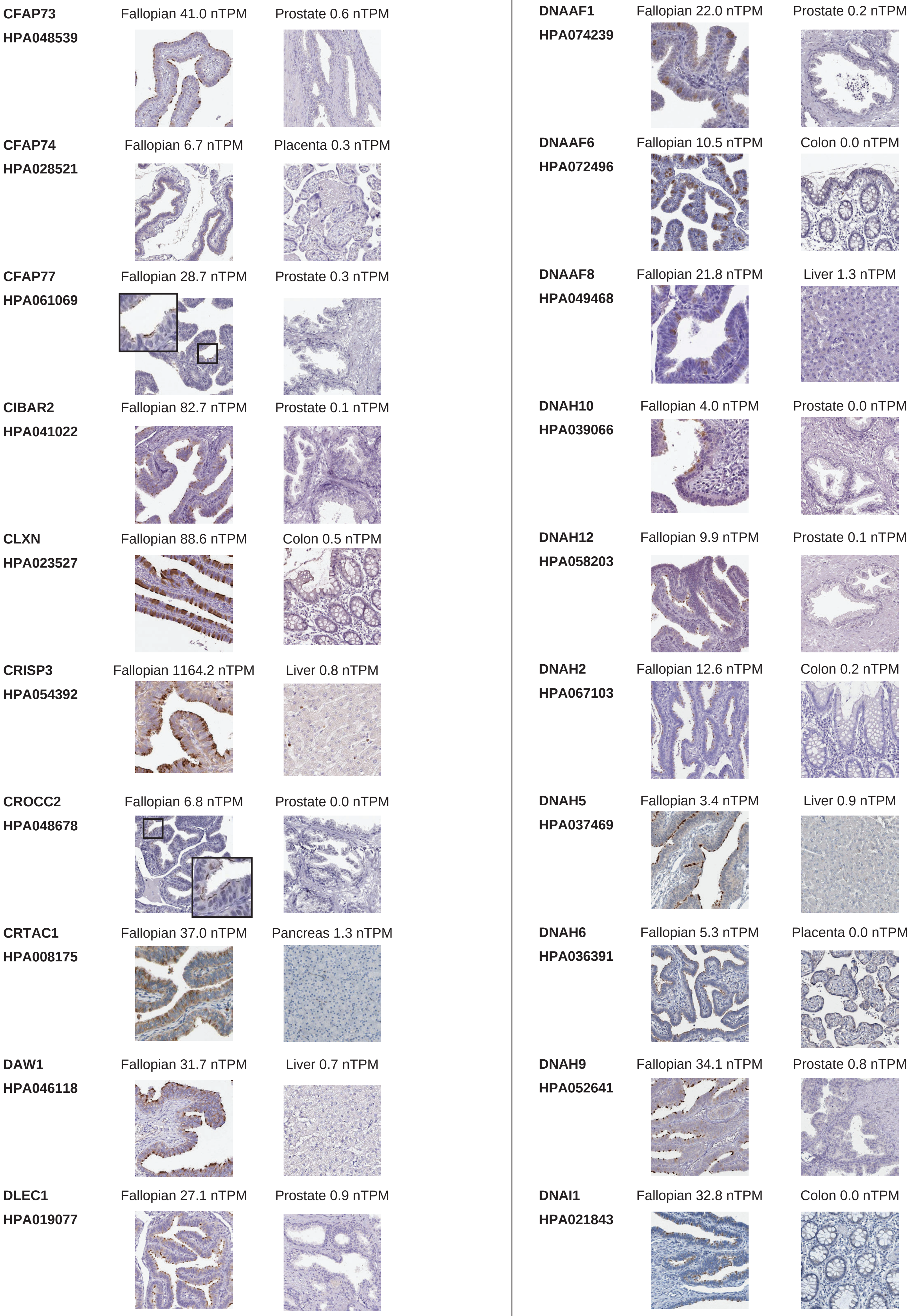
CFAP73
HPA048539
Fallopian 41.0 nTPM
Prostate 0.6 nTPM
CFAP74
HPA028521
Fallopian 6.7 nTPM
Placenta 0.3 nTPM
CFAP77
HPA061069
Fallopian 28.7 nTPM
Prostate 0.3 nTPM
CIBAR2
HPA041022
Fallopian 82.7 nTPM
Prostate 0.1 nTPM
CLXN
HPA023527
Fallopian 88.6 nTPM
Colon 0.5 nTPM
CRISP3
HPA054392
Fallopian 1164.2 nTPM
Liver 0.8 nTPM
CROCC2
HPA048678
Fallopian 6.8 nTPM
Prostate 0.0 nTPM
CRTAC1
HPA008175
Fallopian 37.0 nTPM
Pancreas 1.3 nTPM
DAW1
HPA046118
Fallopian 31.7 nTPM
Liver 0.7 nTPM
DLEC1
HPA019077
Fallopian 27.1 nTPM
Prostate 0.9 nTPM
DNAAF1
HPA074239
Fallopian 22.0 nTPM
Prostate 0.2 nTPM
DNAAF6
HPA072496
Fallopian 10.5 nTPM
Colon 0.0 nTPM
DNAAF8
HPA049468
Fallopian 21.8 nTPM
Liver 1.3 nTPM
DNAH10
HPA039066
Fallopian 4.0 nTPM
Prostate 0.0 nTPM
DNAH12
HPA058203
Fallopian 9.9 nTPM
Prostate 0.1 nTPM
DNAH2
HPA067103
Fallopian 12.6 nTPM
Colon 0.2 nTPM
DNAH5
HPA037469
Fallopian 3.4 nTPM
Liver 0.9 nTPM
DNAH6
HPA036391
Fallopian 5.3 nTPM
Placenta 0.0 nTPM
DNAH9
HPA052641
Fallopian 34.1 nTPM
Prostate 0.8 nTPM
DNAI1
HPA021843
Fallopian 32.8 nTPM
Colon 0.0 nTPM

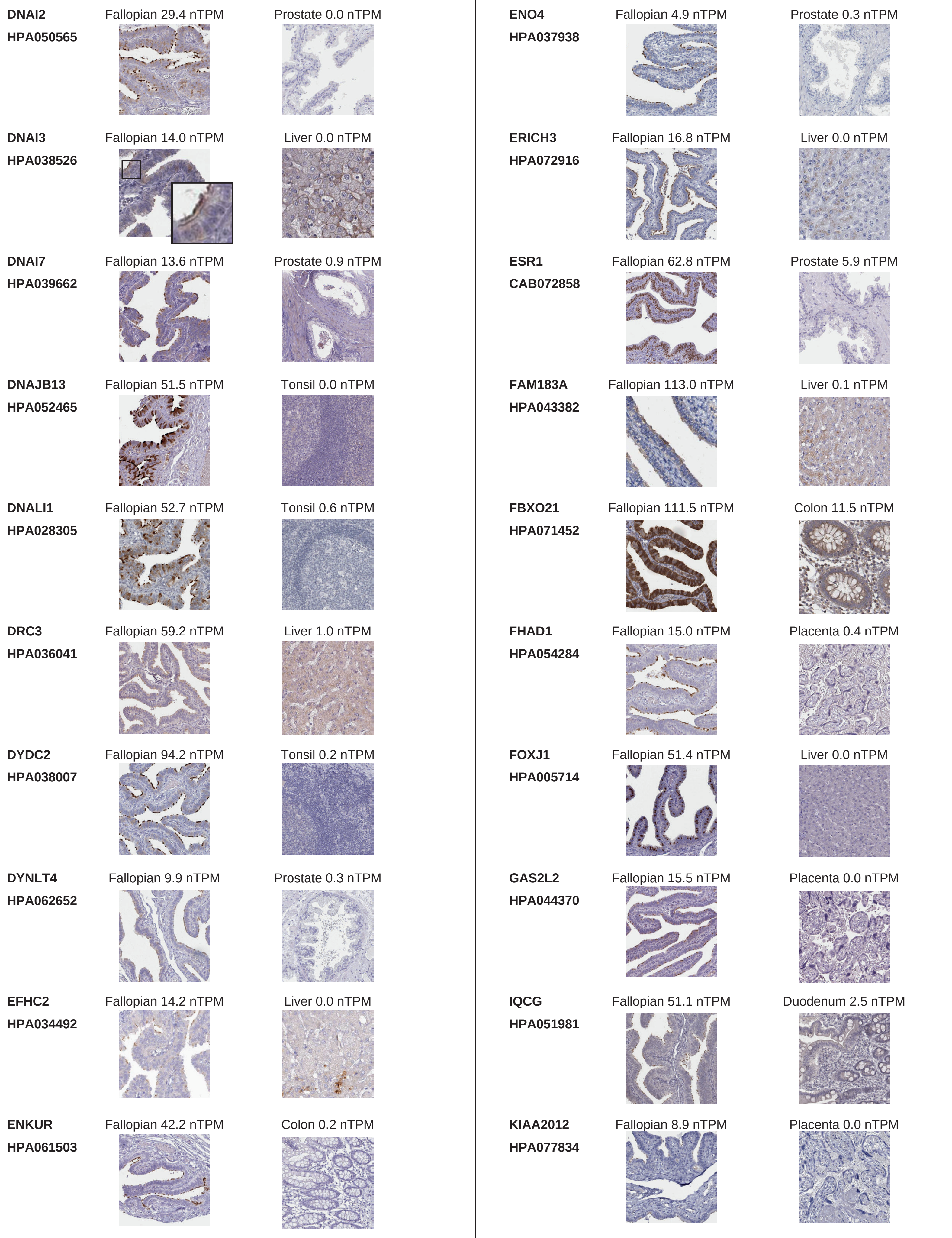
DNAI2
HPA050565
Fallopian 29.4 nTPM
Prostate 0.0 nTPM
DNAI3
HPA038526
Fallopian 14.0 nTPM
Liver 0.0 nTPM
DNAI7
HPA039662
Fallopian 13.6 nTPM
Prostate 0.9 nTPM
DNAJB13
HPA052465
Fallopian 51.5 nTPM
Tonsil 0.0 nTPM
DNALI1
HPA028305
Fallopian 52.7 nTPM
Tonsil 0.6 nTPM
DRC3
HPA036041
Fallopian 59.2 nTPM
Liver 1.0 nTPM
DYDC2
HPA038007
Fallopian 94.2 nTPM
Tonsil 0.2 nTPM
DYNLT4
HPA062652
Fallopian 9.9 nTPM
Prostate 0.3 nTPM
EFHC2
HPA034492
Fallopian 14.2 nTPM
Liver 0.0 nTPM
ENKUR
HPA061503
Fallopian 42.2 nTPM
Colon 0.2 nTPM
ENO4
HPA037938
Fallopian 4.9 nTPM
Prostate 0.3 nTPM
ERICH3
HPA072916
Fallopian 16.8 nTPM
Liver 0.0 nTPM
ESR1
CAB072858
Fallopian 62.8 nTPM
Prostate 5.9 nTPM
FAM183A
HPA043382
Fallopian 113.0 nTPM
Liver 0.1 nTPM
FBXO21
HPA071452
Fallopian 111.5 nTPM
Colon 11.5 nTPM
FHAD1
HPA054284
Fallopian 15.0 nTPM
Placenta 0.4 nTPM
FOXJ1
HPA005714
Fallopian 51.4 nTPM
Liver 0.0 nTPM
GAS2L2
HPA044370
Fallopian 15.5 nTPM
Placenta 0.0 nTPM
IQCG
HPA051981
Fallopian 51.1 nTPM
Duodenum 2.5 nTPM
KIAA2012
HPA077834
Fallopian 8.9 nTPM
Placenta 0.0 nTPM

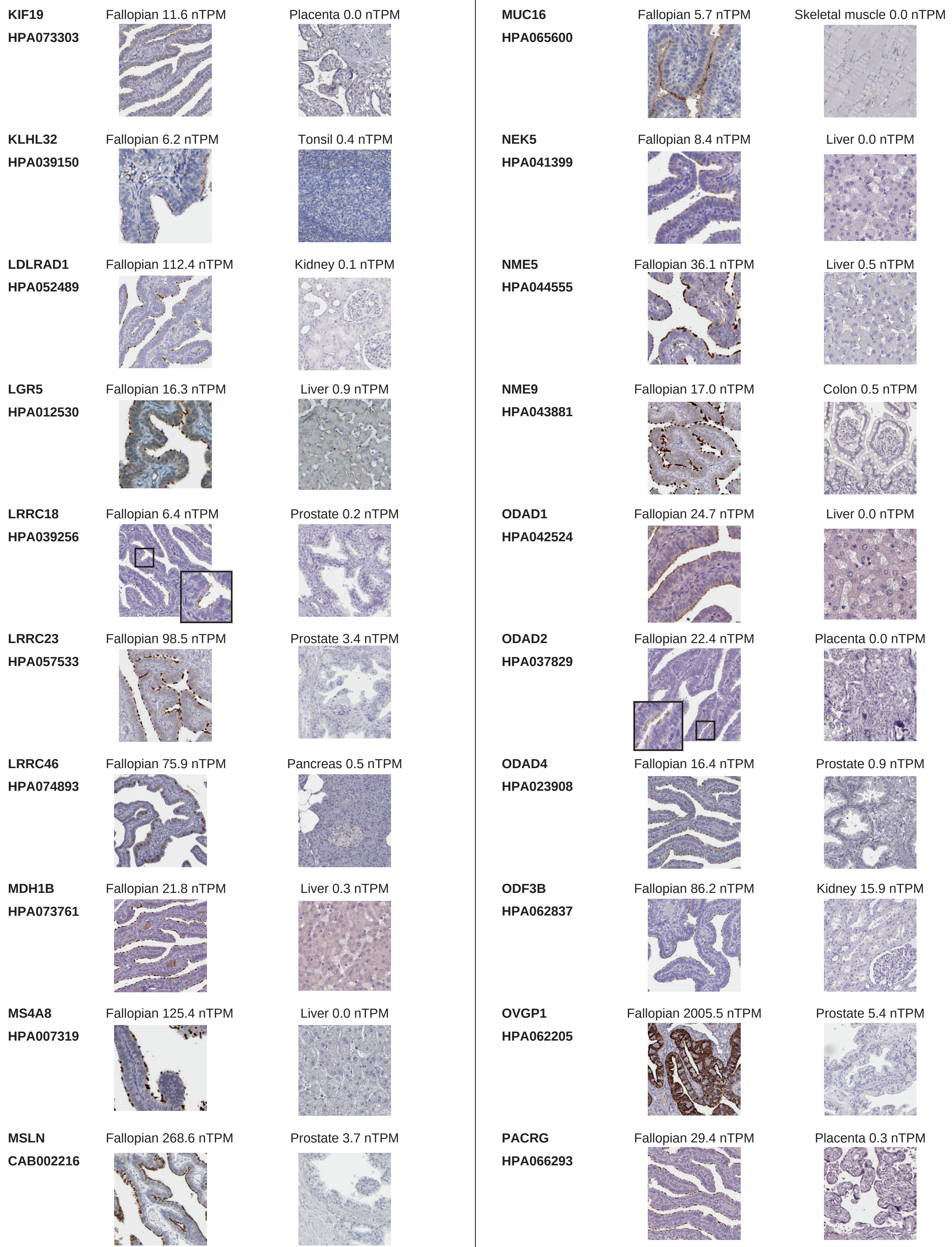
KIF19
HPA073303
Fallopian 11.6 nTPM
Placenta 0.0 nTPM
KLHL32
HPA039150
Fallopian 6.2 nTPM
Tonsil 0.4 nTPM
LDLRAD1
HPA052489
Fallopian 112.4 nTPM
Kidney 0.1 nTPM
LGR5
HPA012530
Fallopian 16.3 nTPM
Liver 0.9 nTPM
LRRC18
HPA039256
Fallopian 6.4 nTPM
Prostate 0.2 nTPM
LRRC23
HPA057533
Fallopian 98.5 nTPM
Prostate 3.4 nTPM
LRRC46
HPA074893
Fallopian 75.9 nTPM
Pancreas 0.5 nTPM
MDH1B
HPA073761
Fallopian 21.8 nTPM
Liver 0.3 nTPM
MS4A8
HPA007319
Fallopian 125.4 nTPM
Liver 0.0 nTPM
MSLN
CAB002216
Fallopian 268.6 nTPM
Prostate 3.7 nTPM
MUC16
HPA065600
Fallopian 5.7 nTPM
Skeletal muscle 0.0 nTPM
NEK5
HPA041399
Fallopian 8.4 nTPM
Liver 0.0 nTPM
NME5
HPA044555
Fallopian 36.1 nTPM
Liver 0.5 nTPM
NME9
HPA043881
Fallopian 17.0 nTPM
Colon 0.5 nTPM
ODAD1
HPA042524
Fallopian 24.7 nTPM
Liver 0.0 nTPM
ODAD2
HPA037829
Fallopian 22.4 nTPM
Placenta 0.0 nTPM
ODAD4
HPA023908
Fallopian 16.4 nTPM
Prostate 0.9 nTPM
ODF3B
HPA062837
Fallopian 86.2 nTPM
Kidney 15.9 nTPM
OVGP1
HPA062205
Fallopian 2005.5 nTPM
Prostate 5.4 nTPM
PACRG
HPA066293
Fallopian 29.4 nTPM
Placenta 0.3 nTPM

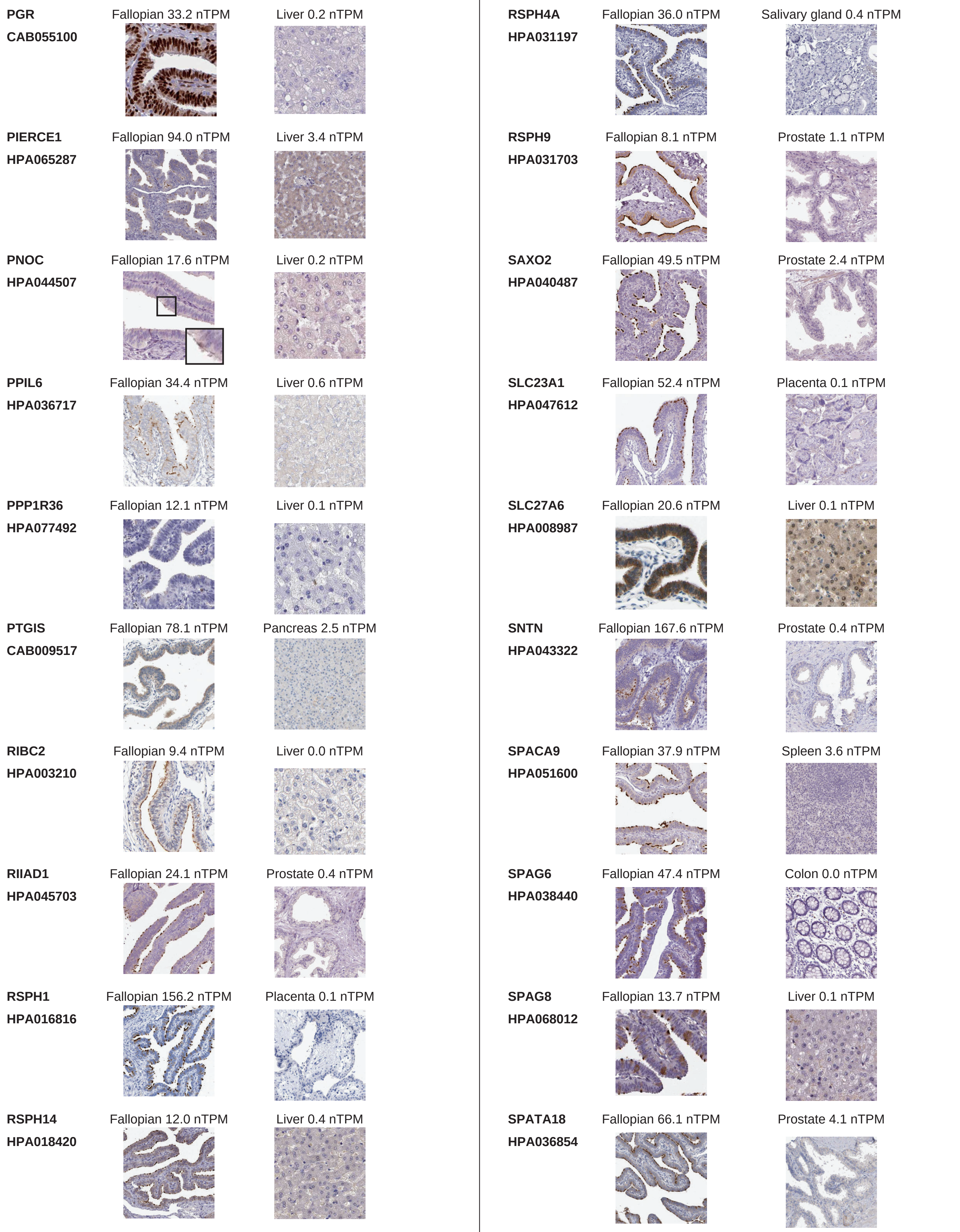
PGR
CAB055100
Fallopian 33.2 nTPM
Liver 0.2 nTPM
PIERCE1
HPA065287
Fallopian 94.0 nTPM
Liver 3.4 nTPM
PNOC
HPA044507
Fallopian 17.6 nTPM
Liver 0.2 nTPM
PPIL6
HPA036717
Fallopian 34.4 nTPM
Liver 0.6 nTPM
PPP1R36
HPA077492
Fallopian 12.1 nTPM
Liver 0.1 nTPM
PTGIS
CAB009517
Fallopian 78.1 nTPM
Pancreas 2.5 nTPM
RIBC2
HPA003210
Fallopian 9.4 nTPM
Liver 0.0 nTPM
RIIAD1
HPA045703
Fallopian 24.1 nTPM
Prostate 0.4 nTPM
RSPH1
HPA016816
Fallopian 156.2 nTPM
Placenta 0.1 nTPM
RSPH14
HPA018420
Fallopian 12.0 nTPM
Liver 0.4 nTPM
RSPH4A
HPA031197
Fallopian 36.0 nTPM
Salivary gland 0.4 nTPM
RSPH9
HPA031703
Fallopian 8.1 nTPM
Prostate 1.1 nTPM
SAXO2
HPA040487
Fallopian 49.5 nTPM
Prostate 2.4 nTPM
SLC23A1
HPA047612
Fallopian 52.4 nTPM
Placenta 0.1 nTPM
SLC27A6
HPA008987
Fallopian 20.6 nTPM
Liver 0.1 nTPM
SNTN
HPA043322
Fallopian 167.6 nTPM
Prostate 0.4 nTPM
SPACA9
HPA051600
Fallopian 37.9 nTPM
Spleen 3.6 nTPM
SPAG6
HPA038440
Fallopian 47.4 nTPM
Colon 0.0 nTPM
SPAG8
HPA068012
Fallopian 13.7 nTPM
Liver 0.1 nTPM
SPATA18
HPA036854
Fallopian 66.1 nTPM
Prostate 4.1 nTPM

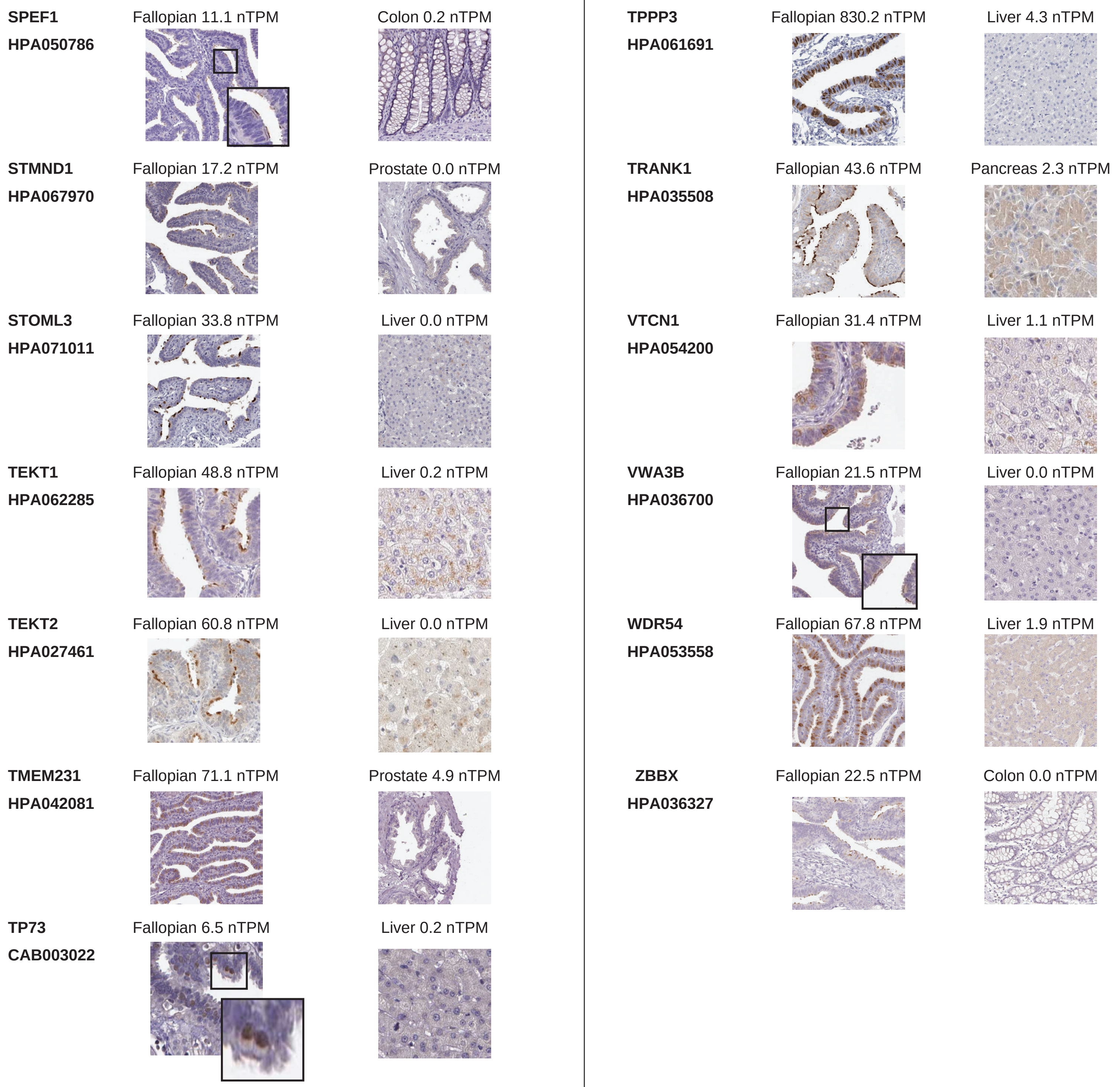
SPEF1
HPA050786
Fallopian 11.1 nTPM
Colon 0.2 nTPM
STMND1
HPA067970
Fallopian 17.2 nTPM
Prostate 0.0 nTPM
STOML3
HPA071011
Fallopian 33.8 nTPM
Liver 0.0 nTPM
TEKT1
HPA062285
Fallopian 48.8 nTPM
Liver 0.2 nTPM
TEKT2
HPA027461
Fallopian 60.8 nTPM
Liver 0.0 nTPM
TMEM231
HPA042081
Fallopian 71.1 nTPM
Prostate 4.9 nTPM
TP73
CAB003022
Fallopian 6.5 nTPM
Liver 0.2 nTPM
TPPP3
HPA061691
Fallopian 830.2 nTPM
Liver 4.3 nTPM
TRANK1
HPA035508
Fallopian 43.6 nTPM
Pancreas 2.3 nTPM
VTCN1
HPA054200
Fallopian 31.4 nTPM
Liver 1.1 nTPM
VWA3B
HPA036700
Fallopian 21.5 nTPM
Liver 0.0 nTPM
WDR54
HPA053558
Fallopian 67.8 nTPM
Liver 1.9 nTPM
ZBBX
HPA036327
Fallopian 22.5 nTPM
Colon 0.0 nTPM

### Supplementary Figure legends

#### Supplementary Figure 1

Heatmap showing the binary relationship of group-enriched tissues (upper) and elevated cell type specificities (lower) for 114 group-enriched genes in FT. The upper heatmap shows which tissues are in the group "group-enriched" for each gene (columns), with a legend showing if the peak RNA level is in FT tissue or not. The lower heatmap shows which cell types are present in the cell type elevated groups for each gene with a legend showing the peak RNA level in Ciliated and/or Secretory cells or not.

#### Supplementary Figure 2

Heatmap showing the binary relationship of enhanced tissues (upper) and elevated cell type specificities (lower) for 117 enhanced genes in FT. The upper heatmap shows which tissues are in the group "enhanced" for each gene (columns), with a legend showing if the peak RNA level is in FT tissue or not. The lower heatmap shows which cell types are present in the cell type elevated groups for each gene with a legend showing the peak RNA level in Ciliated and/or Secretory cells or not.

#### Supplementary Figure 3

Heatmap showing the binary relationship of elevated cell type specificities for 19 genes enriched genes in FT. The bottom legend shows the peak RNA level in Ciliated and/or Secretory cells or not.

#### Supplementary Figure 4

Comparing the list of 123 Ciliated FTE cell-specific proteins to proteomics studies on cilia knowledge databases, ciliopathy databases, and Sperm and flagella datasets. (A) The 123 FT proteins were examined in five different databases for cilia-relevant function and expression. The genes/proteins that are present in any or several resources are labeled in purple or blue. The datasets from Nevers & Karunakaran (blue) are based on predictions and phylogenetic approaches, whereas the others in purple are databases based on literature research. Empty bars represent no data in the databases. List of the 123 proteins examined in IHC compared to (B) datasets of flagellar proteins identified in different studies or (C) a database for ciliopathies called "CiliaMiner" (Turan et al., 2023[34]). The proteins that are present in the dataset are filled in purple or green. The dataset shown in green is based on comparing the proteome of sea urchins, sea anemones, and choanoflagellates[33]; Overlap with this data set provides insight into the conserveness of the proteins we identified. The other flagella protein lists all stem from proteomics studies in human sperm and were collected in Amaral et al., 2014[31]. The data set by Martin-Hidalgo et al[32] lists proteins with changes in phosphorylation profiles related to sperm motility. The comparison to CiliaMiner was conducted with the data present on CiliaMiner's webpage (https://kaplanlab.shinyapps.io/ciliaminer/) on December 03, 2022.

#### Supplementary Figure 5

The expression of FT-elevated proteins in human tissues with motile ciliated epithelial cells. The highlighted proteins represent those stained in both ciliated and non-ciliated cells, and strictly in non-ciliated cells.

**Supplementary Figure 6**

Overview of single-cell clusters in human FT. An UMAP plot showing the different cell clusters identified in human FT by using integration of three scRNA-seq datasets. Each dot represents a cell. The upper plot highlight cells colored by dataset. The bottom represents the cellular identities assigned after manual annotation (see methods for cell identity markers), comprising clusters of ciliated cells, secretory cells, fibroblasts, smooth muscle cells, endothelial cells, immune cells (T cells, B cells, NK cells, and dendritic cells), mast cells, macrophages. The undefined cell cluster was not used in further analysis.

**Supplementary Figure 7**

IHC stainings in human FT by additional antibodies of proteins included in the study.

**Supplementary Figure 8**

(A) IHC staining of FOXJ1 in whole-slide hydrosalpinx (HS) and control FT tissues. (B) Epithelial width measurement for HS and control samples.

**Supplementary Figure 9**

Validation of antibody specificity using positive (FT tissue) and negative control stainings. Negative control tissues were selected based on low mRNA expression, or below detection threshold, and were expected to lack or exhibit very low abundance of the target protein.
